# Insights into the AAV packaging mechanism: Cryo-EM Structure of the AAV2 Rep-Capsid Packaging Complex

**DOI:** 10.64898/2025.12.09.693273

**Authors:** Jason T. Kaelber, Vadim Barnakov, Jiayu Shen, Karen Hernandez, Harrison J. Tarbox, Areeba Khan, Carlos R. Escalante

**Author notes:** To whom correspondence should be addressed. Correspondence may also be addressed to Jason Kaelber.

## Abstract

The Adeno-associated virus (AAV) has become the most used viral vector for gene therapy applications to treat monogenic diseases, with over 250 clinical trials and six FDA-approved biologics. AAV has a single-stranded DNA genome encapsulated in a 60-subunit icosahedral protein shell. Assembly of empty capsids occurs in the nucleus, and in a subsequent step, the genome is packaged by the motor activity of the AAV Rep proteins. The translocation of ssDNA has been suggested to occur through one of the twelve channels at the fivefold symmetry axis. While there is substantial evidence that Rep proteins directly interact with the capsid, the specific molecular determinants, stoichiometry, and translocation mechanism remain unknown. To understand how Rep proteins assemble on the capsid, we examined Rep-capsid complexes using cryo-electron microscopy with single-particle reconstruction. Our results show that Rep proteins can assemble into the capsid fivefold pore as either pentameric or hexameric rings. The different ring complexes dock into the pore similarly, using the post-sensor1 β-hairpin motif (pos1βh2) as a capsid interaction module. This interaction induces the folding of a segment in the capsid HI loop, resulting in an expansion of the pos1βh2 β-sheet. Our structures show that any of the AAV Rep proteins can interact with the capsid, and this mode of interaction may be a conserved mechanism across all parvoviruses. Collectively, our results provide insights into the mechanism of AAV genome encapsidation and could inform strategies to improve recombinant AAV packaging efficiency.

## Introduction

Adeno-associated virus (AAV) is a small, non-enveloped, single-stranded DNA (ssDNA) parvovirus that is currently the leading platform for gene therapy vector development^1–4^. AAV’s appeal lies in its non-pathogenic nature, relatively low immunogenicity, broad tissue tropism, and ability to infect both dividing and non-dividing cells^5–7^. Based on their genome size and the limited number of proteins they encode, AAVs are among the simplest viruses in nature. Their ssDNA genome is approximately 4.7 kb and contains only two genes: the *rep* gene and the *cap* gene. The *rep* gene encodes four nonstructural Rep proteins—Rep78, Rep68, Rep52, and Rep40—named according to their apparent molecular weights. The *cap* gene produces three viral proteins (VPs): VP1, VP2, and VP3, as well as a small assembly factor AAP^8^ and the membrane-associated accessory protein MAAP^9^. The simplicity of the parvovirus genome demands that its non-structural Rep proteins be multifunctional, managing critical functions such as DNA replication, gene expression regulation, and genome packaging. The AAV capsid contains 60 copies of VP1-3 proteins in a 1:1:10 ratio, arranged in a *T*=1 icosahedral symmetry, forming a quasi-spherical particle about 26 nm^10,11^. A prominent structural feature on the capsid surface is a cylindrical channel or pore located at the five-fold symmetry axis, formed by five DE β-hairpins, believed to be the site of genome packaging^12^. In AAV and most parvoviruses (with the exception of certain densoviruses that reverse these dynamics), the five-fold pore is mostly empty before genome packaging, but becomes filled with the N-terminus of the capsid protein after packaging.^13^

AAV’s ideal status as a gene therapy vector stems from its unique properties, including its ability to effectively transduce both dividing and non-dividing cells, relatively low immunogenicity, and broad tissue tropism. Due to these advantageous properties, there are currently seven FDA/EMA-approved biologics for the treatment of monogenic disorders, including lipoprotein lipase deficiency, hemophilia B, retinal dystrophy, and Duchenne muscular dystrophy^14^. Despite the successful implementation of AAV vectors, a persistent issue in the production of recombinant AAV (rAAV) is the abundance of empty and partially filled capsids^15–19^. Depending on the expression and purification systems used during rAAV production, between 70 and 90% of empty capsids are generated, necessitating a multistep purification process that increases manufacturing costs^20,21^. Understanding the AAV packaging mechanism can help overcome these challenges.

Over the past 40 years, various research groups have delineated the general steps leading to the formation of infectious AAV particles. Assembly of AAV capsids occurs in the nucleus and only requires the presence of VP3 and the small assembly protein AAP, which promotes VP stability and assembly^8,22–25^. In contrast to the papillomaviruses and polyomaviruses (in which capsids condense around a genome), capsid assembly and genome packaging occur sequentially, as ssDNA is inserted into pre-formed empty capsids.^26^ In the absence of the non-structural AAV Rep proteins, the empty capsids accumulate in the nucleoli but move to the nucleoplasm in a Rep-dependent process^23,27^. Coimmunoprecipitation studies have shown that all four Rep proteins are associated with the capsid; however, the identity of the final complex has not been determined^28,29^. In 2001, King et al. demonstrated that packaging is mediated by the helicase activity of the Rep proteins. They showed that eliminating Rep40/Rep52 expression resulted in a 20-fold reduction in total encapsidation, a significant decrease in the amount of full-length genome inserted, and a 200-fold reduction in infectious particle titer. Moreover, they demonstrated that DNA encapsidation proceeds in a 3’ to 5’ direction that parallels the DNA translocation direction of the Rep proteins^30^. Subsequent studies indicate that lysine residues in the Reps pre-sensor 1 β-hairpin (PS1βH) motif are necessary for efficient packaging^31^. Bleker et al. conducted a study showing that mutations around the fivefold pore of AAV capsids hinder genome packaging. This impact seems to result from steric hindrance at the pore. Similarly, research on the minute virus of mice (MVM) showed that a leucine-to-tryptophan mutation in the MVM capsid’s five-fold pore also caused near-total disruption of packaging, indicating that the 5-fold pore is the DNA packaging portal^12,32,33^. The sequential packaging systems of tailed bacteriophages and herpesviruses are among the most extensively studied viral packaging systems, sharing similar structural and functional features for packaging dsDNA^34–43^. These systems have a dedicated vertex for packaging, with a portal protein that serves as a platform for assembling the other components of the packaging motor^44^. By contrast, in adenoviruses^45^ and parvoviruses, no portal has been found, raising the question of how the symmetrical capsid selects a single unit-length genome for encapsidation. Beyond preexisting capsid asymmetry (such as a portal), binding of a receptor to a single vertex of a perfectly icosahedral capsid can induce asymmetry and activate that site alone^46,47^, so by analogy, it is conceivable that attachment of Rep to one capsid pore could induce a change that would preclude other Rep from binding at the other 11 pores. Alternatively, if Rep can bind multiple vertices simultaneously, another mechanism must be at play to prevent DNA collisions.

Because it is possible to carry out a packaging reaction with AAV *in vitro*,^48,49^ we reasoned that it should be feasible to capture a physiologically-relevant complex between Rep proteins and the capsid by crosslinking. To elucidate how AAV Rep proteins interact with the capsid, we use cryo-electron microscopy to determine the structure of the Rep helicase domain in complex with empty AAV2 capsids. Here, we show that Rep proteins can assemble into the capsid fivefold pore as pentameric or hexameric rings. The structures dock into the pore, similarly, using the post-sensor1 β-hairpin motif (pos1βH2) as a capsid interaction module. The interaction induces folding of a region of the capsid HI loop, leading to an expansion of the pos1βH2 β-sheet. Our structures show that any of the AAV Rep proteins can interact with the capsid, and that this interaction mode may be a conserved mechanism across all parvoviruses. Taken together, our results provide insights into the mechanism of AAV genome encapsidation and may guide strategies to enhance recombinant AAV packaging efficiency.

## Materials and Methods

### Expression and Purification of Rep proteins

Rep68 was expressed in *E. coli* BL21(DE3) cells (Novagen) using a modified pET-15b vector containing a HRV-3C protease site. All proteins were purified as previously described ^50^. For structural studies, a Rep68 containing a C151S mutant will be referred to as Rep68 in the manuscript ^51^. The final buffer contained 25 mM Tris-HCl pH 8.0, 200 mM NaCl, and 2 mM TCEP. The HRV3C protease was expressed in BL21(DE3)-pLysS at 37 °C for 3 hours in LB medium containing 1 mM IPTG. Cell pellets were lysed in Ni-Buffer A (20 mM Tris-HCl pH 7.9 at 4 °C, 500 mM NaCl, 5 mM imidazole, 10% glycerol, and 1 mM TCEP). After five 10-s cycles of sonication, the fusion protein was purified using a Ni-column equilibrated in Ni-buffer A. Protein eluted was desalted using buffer A and a HiPrep^TM^ 26/10 desalting column (GE Healthcare). The hexahistidine tag was removed by treating the protein with HRV3C protease (150 μg protease per mg His-PP-Rep68) with overnight incubation at 4 °C. A second Ni-column chromatography removed the uncleaved fusion protein and untagged Rep68. Rep68 was finally purified by gel filtration chromatography using a HiLoad Superdex 200 16/600 PG column (GE Healthcare) and Size Exclusion buffer (25 mM Tris pH 8.0, 200 mM NaCl, 1 mM TCEP). N-terminus His6-tagged WT and mutant Rep68 proteins were concentrated to 2 mg/ml with 50% glycerol, flash-frozen in liquid N_2_, and stored at −80°C.

Rep210 (residues 210-536) was also expressed in E. coli BL21(DE3) at 18°C using a modified pET-15b plasmid with tobacco Etch virus (TEV) protease site and purified as described before^52^. The final buffer contains (25 mM Tris-HCl [pH 8.0], 200 mM NaCl, and 2mM TCEP).

Rep40 was expressed as described previously^53^. In short, plasmid pET-15b with a tobacco Etch Virus (TEV) protease site was used, and expression was done at 18°C. Subsequent steps were similar to Rep68 purification.

### Preparation of Rep-Capsid Complexes

Rep68, Rep210, and Rep40 were reconstituted in 1x PBS (pH 7.4) using a Hitrap^™^ desalting column (Cytiva). Protein concentrations were determined by measuring absorbance at 280 nm and using the calculated extinction coefficient. Rep proteins were mixed with ssDNA and incubated on ice for 10 minutes. Rep210 was mixed with ssDNA at a 6:1 molar ratio. Subsequently, ATPγS and MgCl₂ were added for final concentrations of 5 mM. Empty AAV2 capsids (5×10^12^ total particles) purchased from AAVnerGene were mixed with the Rep210-ssDNA complex and incubated on ice for 5 minutes at a molar ratio of 1 VP3 protein to 36 Rep210 (corresponding to 2160 rep proteins per capsid). The complex was concentrated to a final volume of 50 µL and cross-linked with bis(sulfosuccinimidyl)suberate (BS^3^) by incubating on ice for 90 minutes. Crosslinking was quenched by adding 30 mM ammonium bicarbonate for 15 minutes. For the Rep68, a Rep68-ITR complex was mixed with the capsid at a molar ratio of 1 VP3 to 80 Rep68 complex. Crosslinking was performed as previously described. Samples were stored at 4°C until grid preparation.

### Imaging and icosahedral reconstruction of the Rep210-Capsid complex

13,369 micrograph movies were acquired by EPU software using a K3 detector on a 300kV “Krios G3i” TEM with BioQuantum energy filter set to 10 eV slit width. Field-standard parameters were used except that: an objective aperture of 100μm diameter was used, and the total dose was 73.15*e*^-^/Å^2^. The increased dose was chosen to accumulate a low-resolution signal for pseudosymmetry breaking.^54^ The nominal pixel size of 0.813 Å/pixel was calibrated after reconstruction to 0.82 Å/pixel by comparison of the capsid to the crystal structure of AAV2.

Micrographs were processed for icosahedral reconstruction in cryoSPARC v. 4.5.3 by: patch motion and CTF correction; exclusion of 23% of micrographs due to CTF fits >4 Å or bad ice; template-based picking; 2D classification to exclude “bad” particles; separation of micrographs into beam-shift groups by parsing of the XML metadata files from EPU; and refinement with per-particle defocus fitting, per-group fitting of aberrations up to the fourth order, per-group fitting of anisotropic magnification, and Ewald sphere correction. After correcting the handedness of the reconstruction, an icosahedrally-symmetrized map of 145,050 capsids at 1.83 Å was obtained.

### Rep210-Capsid structure determination

To confirm Rep binding at the fivefold vertex and understand the location and extent of rep, asymmetric local refinement was performed using various mask and fulcrum positions. This resulted in a pseudosymmetric medium-resolution map that shows rep shapes in a pentameric configuration at the five-fold vertex. This does not prove that rep is a pentamer because of the strong pseudosymmetry bias from icosahedral reconstruction,^54^ but it helped generate masks in EMAN2 for particle classification. Capsid-subtracted particles were symmetry-expanded and a soft mask (around the rep region but excluding any capsid volume) was applied for three rounds of 3D classification at 12 Å filter resolution. Class grouping/splitting was performed manually based on inspection of the volumes. We noted that some classes appeared virtually identical to other volumes if a rotation of a multiple of 72° was applied, as would be expected if an asymmetric complex docks on a five-fold symmetrical vertex and sorting—but not realignment—of symmetry-expanded copies of each particle is permitted.

Only in the final local refinement was any rotatational/translational deviation from the symmetry-expanded transform permitted. This prevented leakage of one symmetry-expanded state into another icosahedrally-related state, which could have resulted in constructive interference of noise, overestimation of resolution, and loss of some classes. Because the mask used for classification and local refinement excluded parts of rep at the capsid-interacting interfaces, and because capsid-subtracted particles were used, the transforms for each locally-refined particle were mapped back onto the non-subtracted particles and reconstructed to create a map containing both rep and cap together. These maps are of marginally lower resolution than the maps generated from capsid-subtracted particles and both were used in tandem for molecular modeling and can be accessed from the databank under the “additional maps” download option.

Modeling was performed by iterating between manual building with coot and molecular dynamics flexible fitting with ISOLDE^55^ starting from initial models of the capsid (AAV2 L336C, PDB 6E9D) and rep40:ADP (PDB 1U0J). Deviations from rotational symmetry were quantified in ChimeraX^56^ and can be recapitulated with four copies of the structure open using the following command: delete #1-4/A-E #1-2:1-277 #3-4:278-999; changechains #2 F,G,H,I,J,K I,J,K,F,G,H; changechains #4 F,G,H,I,J,K I,J,K,F,G,H ; align #1 to #2; align #3 to #4.

### Imaging and reconstruction of Rep68-capsid complex

6,634 micrograph movies were acquired as for Rep210, but with a nominal pixel size of 1.1Å. 3,081 micrographs were included with CTF <8.85 Å, good ice, and at least one particle present, and a map of 2.25 Å was reconstructed from 7,760 capsids.

Masks for local refinement were recycled from Rep210 reconstruction and an analogous workflow was used for masked classification and focused refinement.

### 2D reconstruction of Rep oligomers bound to capsids

The capsid-subtracted particles used as inputs for Rep210/cap or Rep68/cap 3D classification were taken—without symmetry expansion—as inputs for 2D classification by CryoSPARC version 4.5.3. For assessment of the occupancy of multiple vertices simultaneously, a soft circular mask with approximate inner diameter of 415 Å was used, whereas for assessment of the oligomeric states present at a single vertex, a soft circular mask with diameter of 148 Å was used; in both cases, the initial classification uncertainty factor was increased from default to 4 and the maximum alignment resolution was truncated at 8 Å to avoid overfitting.

### 2D reconstruction of Rep oligomers not bound to capsids

For micrographs of Rep210/cap and Rep68/cap (separately), Rep oligomers not associated with capsid particles were also reconstructed. For Rep210, parameters for boxing using an elliptical template were optimized from manual boxing of a small number of micrographs. 2D classification of these “blob picks” yielded 2D classes of adequate shapeliness for template-based boxing, which produced an improved set of particles whose 2D classes were more featureful. Multiple oligomeric states were best visualized when the CryoSPARC software parameter “Initial classification uncertainty” was tuned to 1.0 or lower, but there was no difference in the presence/absence of states between the initial blob pick and later template pick classes.

### ATPase Assay

A direct colorimetric assay was used to determine ATPase activity by measuring the released inorganic phosphate during ATP hydrolysis using the malachite green method^57^. Reaction conditions were modified using the instructions of Malachite green phosphate assay kit (Sigma-Aldrich MAK307-1KT). Protein concentration was kept at 60 nM, and the reaction was performed at 30° for 30 min on 96-well plates. Results were reported as the amount of released inorganic phosphate (Pi).

### Helicase Assay

The substrate used in this assay is a heteroduplex DNA consisting of an 18-bp duplex region with a 15-nucleotide 3’ tail at the bottom strand. The top strand has the sequence 5’- AGAGTACGGTAGGATATGAACCAGACACATGAT-3’; the bottom strand with sequence 5’-CATATCCTACCGTACTCT-F-3’ is labelled at the 3’ end with fluorescein and is released upon unwinding. We used the unlabeled bottom strand as a trap to prevent the reannealing of the displaced fluorescent strand. All reactions were performed in a buffer containing 25 mM HEPES, 50 mM NaCl (pH 7.0) in a total volume of 50μl. Rep68 at different concentrations was mixed with double-stranded F-DNA at a final concentration of 1 μM and incubated for 15 min. The reaction was started by adding 5mM ATP-Mg and 2.5 μM trap DNA. The reaction was incubated at 25°C for 1 minute. EDTA was used to stop the reaction at a final concentration of 20μM. Aliquots of 10μl were loaded in a 12% bis-acrylamide gel (30%) (19:1) using 6X-loading dye (0.25 xylene cyanol FF, 30% glycerol). A Gel Doc EZ Imager was used for densitometry and band analysis. Background lane subtraction, white illumination, and an activation time of 300 sec were used for the analysis.

## Results

### Structure determination of AAV2 Capsid-Rep complexes Rep210-Capsid complex

The CryoEM micrographs and 2D class-average images clearly showed AAV2 particles decorated with Rep molecules. Icosahedral reconstruction yielded a symmetrized map with a resolution of 1.83 Å (Fig. 1A). Further inspection showed that the appearance of holes within the rings of aromatic residues is consistent with the computed resolution (Supplementary Figure 1). Rep density was present in the symmetrized map, albeit blurred and at lower occupancy than the capsid density, indicating misalignment and/or heterogeneous composition.^54^ Capsid-subtracted particles were symmetry-expanded, and the area around a single vertex was classified. The most common class in the initial 3D classification was Rep-free vertices (23%), but combined, the classes with six copies of Rep constituted the majority of vertices. Classes containing pentameric Rep were also observed. Local refinement of a hexameric class representing 14.6% of the expanded particles resulted in a nominal resolution of 2.9 Å with excellent density for a subset of the Rep210 subunits (Fig. 1B), but density for other subunits was weak and contained “ghost” densities characteristic of superposition of multiple conformations, indicating that further classification was required. After iterative rounds of classification, two conformations of hexameric Rep were isolated (Fig. 1D-E), with each reconstruction representing about 3% of the initial data, and one conformation of pentameric Rep representing about 0.6% of the initial data (Fig. 1C). These frequencies do not reflect the prevalence of these configurations in solution.^58^ Despite the lower nominal resolutions of 3.09 Å and 3.00 Å for the hexameric conformations, these maps enabled more complete modeling of all six rep subunits and exhibited fewer artefacts than the superset.

**Figure 1.**
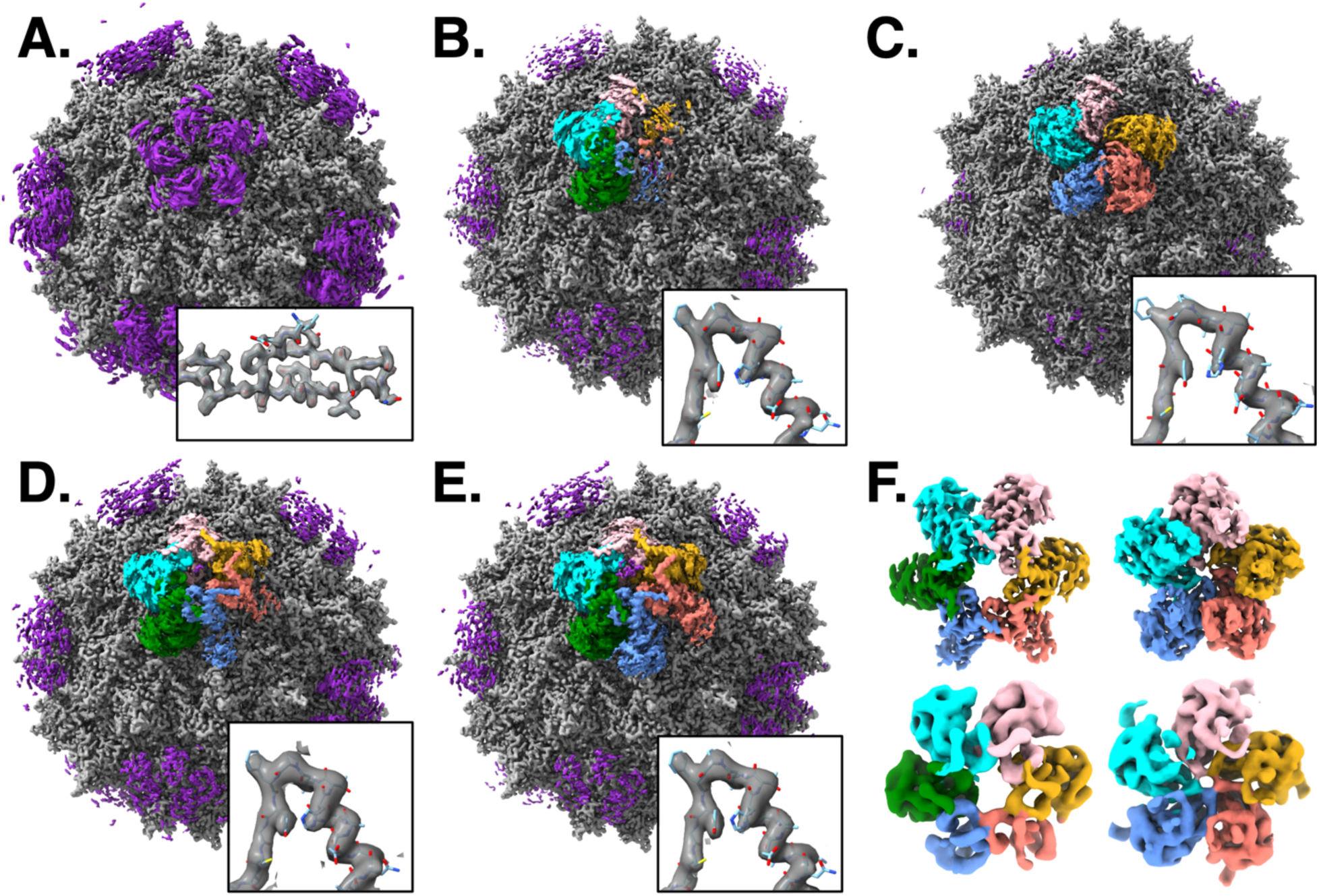
Structures of Rep/Capsid complexes. (A) Standard icosahedral reconstruction of rep210/AAV2 shows clear density for the capsid (gray) as well as non-capsid density (purple). Inset zooms in on capsid VR-IX (residues 710-730), transparent with model displayed. (B) Initial asymmetric local refinement at one vertex revealed additional molecular details of rep structure but did not accurately reflect the oligomeric state. To visualize the Rep210 oligomer in context, refined orientations were mapped back from capsid-subtracted particles onto non-capsid-subtracted particles. These non-subtracted maps are shown in panels B-E. Inset for panels B-E zooms in on Rep α8-β2 (residues 340-358) in the capsid-subtracted maps of Rep210 alone. (C) Asymmetric local refinement at one vertex of the class of particles containing pentameric Rep210 representing 0.6% of all particles. (D) Asymmetric local refinement at one vertex of a class of particles (“complex 1”) containing hexameric Rep210 representing ∼3% of all particles. (E) As panel D, but the class representing complex 2. (F) Focused reconstructions of the rep at one vertex based on capsid-subtracted particles. Representative hexameric and pentameric classes are shown for each of rep210 (top) and rep68 (bottom).

### Rep68-Capsid Complex

Because of the large excess of Rep68 relative to capsid in this experiment, micrographs largely consisted of close-packed Rep68 oligomers with an average of 2.3 capsids per micrograph. However, when those 7,760 capsids were averaged in 2D or 3D, a clear Rep68 density was observed at identical sites to those of the Capsid-Rep210 complexes. After classification of capsid-subtracted particles, local refinement of 45,150 symmetry-expanded particles yielded a hexameric conformation of Rep68 on the capsid with nominal resolution of 6.1 Å. In comparison, 75,559 particles contributed to a pentameric conformation with nominal resolution of 5.5 Å (Fig. 1F, bottom). No nontrivial differences were noted between the Rep68 and Rep210 maps at the moderate resolutions achieved (Fig. 1F), except for near the N-terminal end of resolved density. Therefore, Rep210 maps were interpreted in more detail to understand the atomic details of the interaction.

### Structure determination of AAV2 Rep210-Capsid complexes

The CryoEM micrographs and 2D class-average images clearly showed AAV2 particles decorated with Rep molecules (Figure 2A). The maps showed that Rep density varied across the particle surface, suggesting different degrees of occupancy. The most populous class in the initial 3D classification was Rep-free vertices (23%), but together, classes containing 6 copies of Rep accounted for the majority of vertices. Classes containing pentameric Rep were also present. Local refinement of a hexameric class representing 14.6% of the expanded particles resulted in a nominal resolution of 2.9Å with excellent density for a subset of the Rep210 subunits, but density for other subunits was weak and contained “ghost” densities characteristic of superposition of multiple conformations,^54^ indicating that further classification was required. After iterative rounds of classification, two conformations of hexameric Rep were isolated, with each reconstruction representing about 3% of the initial data, and one conformation of pentameric Rep representing about 0.6% of the initial data. These frequencies do not reflect the prevalence of these configurations in solution. Despite the lower nominal resolutions of 3.09 and 3.00 Å for the hexameric conformations, these maps enabled more complete modeling of all six rep subunits and exhibited fewer artefacts than the superset.

**Figure 2.**
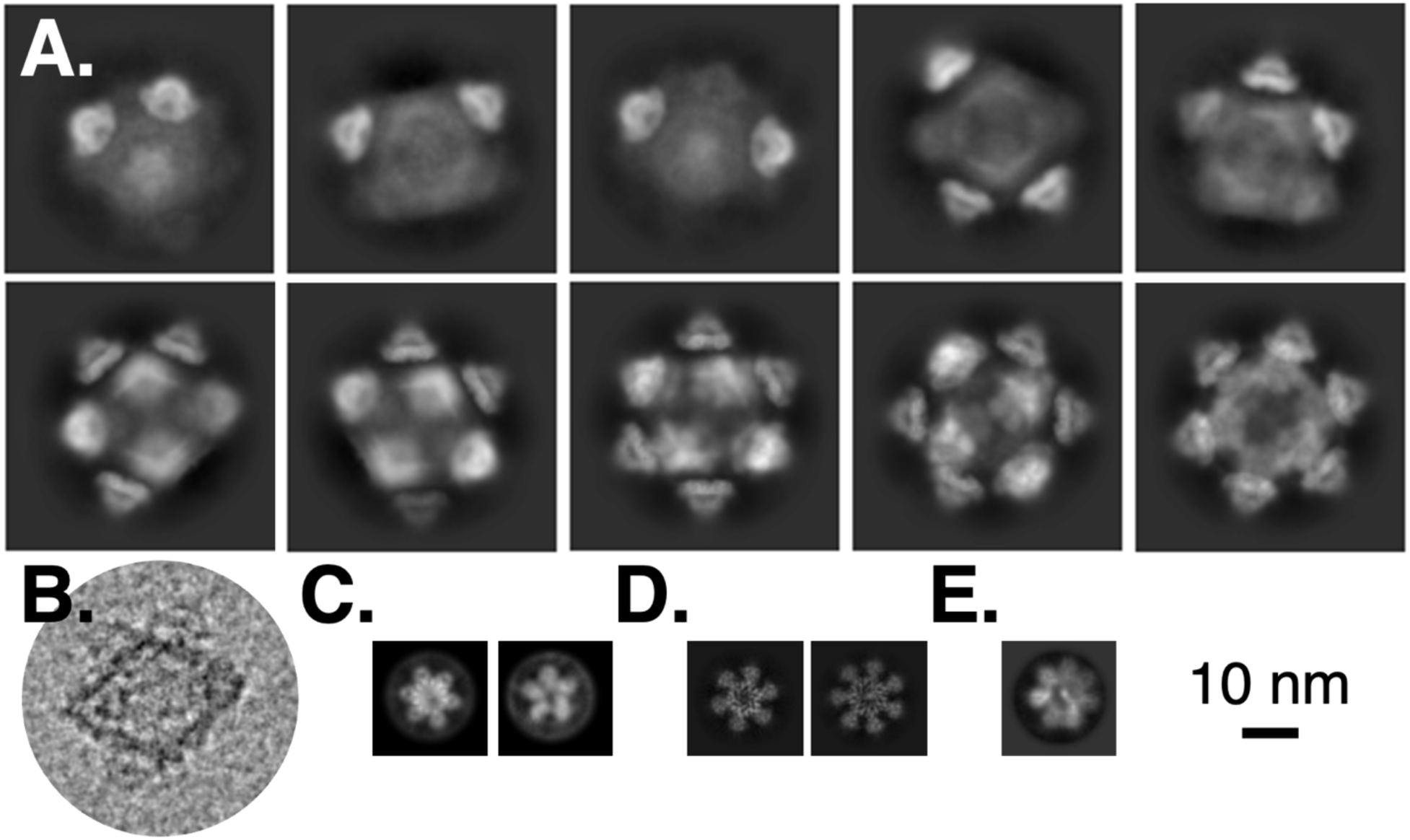
2D characterization of Rep oligomers at vertices. (A) 2D class-averages of capsid-subtracted rep210/capsid particles show diversity in the number of bound rep complexes and their relative configuration. (B) In a representative particle (filtered to 2 nm resolution) of rep210/capsid, rep-occupied and rep-unoccupied vertices can be seen. (C) 2D class-averages of capsid-subtracted rep210/capsid particles with a tight mask around the central region (to eliminate coupling of orientations between vertices) reveal hexameric and pentameric oligomers. (D) 2D class-averages of free rep210 oligomers (extracted from micrographs that also contain rep210/capsid complexes) reveal hexameric and heptameric oligomers. (E) 2D class-averages of free rep68 oligomers (extracted from micrographs that also contain rep68/capsid complexes) reveal heptameric oligomers. Same scale bar for each panel.

### Vertex occupancy

We queried whether Rep oligomers simultaneously occupy one or many vertices. When we mix Rep proteins with preformed empty capsids to mimic the molecular interactions that occur in natural infections, we observe multiple vertices per capsid decorated with Rep complexes. In fact, binding at multiple vertices is seen in raw particle images (Fig. 2A), 2D class averages (Fig. 2B), and 3D classifications of vertices (Fig. 1F). When aligning Rep orientation based on one vertex, the density at other vertices appears smeared and follows the fivefold icosahedral pseudosymmetry (Fig. 1B-E), indicating a lack of conformational coupling between neighboring vertices. Furthermore, we note that Rep binding to a single vertex on AAV capsids does not cause conformational changes capable of transmitting information to a distant site. However, a caveat is that our structures lack ssDNA, which could induce structural changes.

### Structures of Capsid-Rep Complexes

Parvoviral AAV Rep proteins exist as several isoforms, and within an isoform, the protein can adopt multiple oligomeric states. The large Rep proteins Rep68/Rep78 exist as a mixture of oligomers, ranging from dimers to octamers^51,59^. In contrast, the small Reps, Rep40/Rep52, are monomeric but can form transient dimers upon ATP binding^52,60^. The Rep210 used in this study contains the complete Rep68 helicase domain, which includes the linker latch motif (residues 210-224) and Rep40 (residues 225-536). Thus, Rep210 resembles Rep40 in lacking the OBD but also resembles Rep68 in its ability to oligomerize^52^. When Rep210 binds ssDNA, it can form heptameric or hexameric rings, with the latter becoming predominant in the presence of ATPγS. Although free Rep210 and Rep68 heptamers were observed in the grids (Figure 2E-F), they were not found bound to the capsid. Conversely, hexamers are found both in the unbound state and bound to the capsid (Figure 2C-D). We do not observe any other free Rep oligomeric forms, but surprisingly, a significant minority of capsid-bound Rep210 exists as a pentamer (Figure 2C).

### Pentameric Complex

Reconstructing Rep-Cap complexes generally will produce maps of the pentameric Rep unless careful measures are taken to avoid pseudosymmetry artifacts resulting from the capsid’s strong icosahedral symmetry in scattering. One way we increased our confidence that the pentameric capsid-bound state was not an artifact was by running 2D class-averaging of capsid-subtracted particles (Figure 2C). Prior orientation information is not present in the 2D classification process, and the contribution of the capsid is lessened; it is unlikely (albeit conceivable) that the residual from capsid subtraction drives correlations in the class-average because no features aside from copies of Rep can be seen. Additionally, the 3D reconstruction of the pentameric Rep210-Capsid complex exhibits asymmetry within the pentamer (Figure 1C, F), including biochemically reasonable differences between subunits that are not explained by pseudosymmetry overlaps.

The five Rep molecules form a ring-like structure measuring 100 Å × 70 Å × 100 Å. The ring is asymmetric, divided into trimeric and dimeric halves with the Rep molecules oriented so that their pre-sensor-1 β-hairpins (ps1βh) face the ring’s center (Figure 3A). The pentameric arrangement imposes geometric constraints on the size of the Rep ring’s central cavity, which is closed at the ps1βh level, preventing ssDNA from passing through (Figure 3B). The arrangement of the individual Rep subunits is established by interactions with the pore spikes, which keep them sufficiently apart to prevent any interactions between the OD domains, giving the assembly a Rep40-like behavior (Figure 3C). The absence of OD-OD interaction results in weak interfaces between subunits, with the buried surface area (BSA) not exceeding 849 Å², compared to approximately 2100 Å² for the Rep68 dimer interface when ATPγS is bound^61^ (Figure 3D). Even though grids were prepared with ssDNA and ATPγS, the map shows no clear densities indicative of their presence.

**Figure 3.**
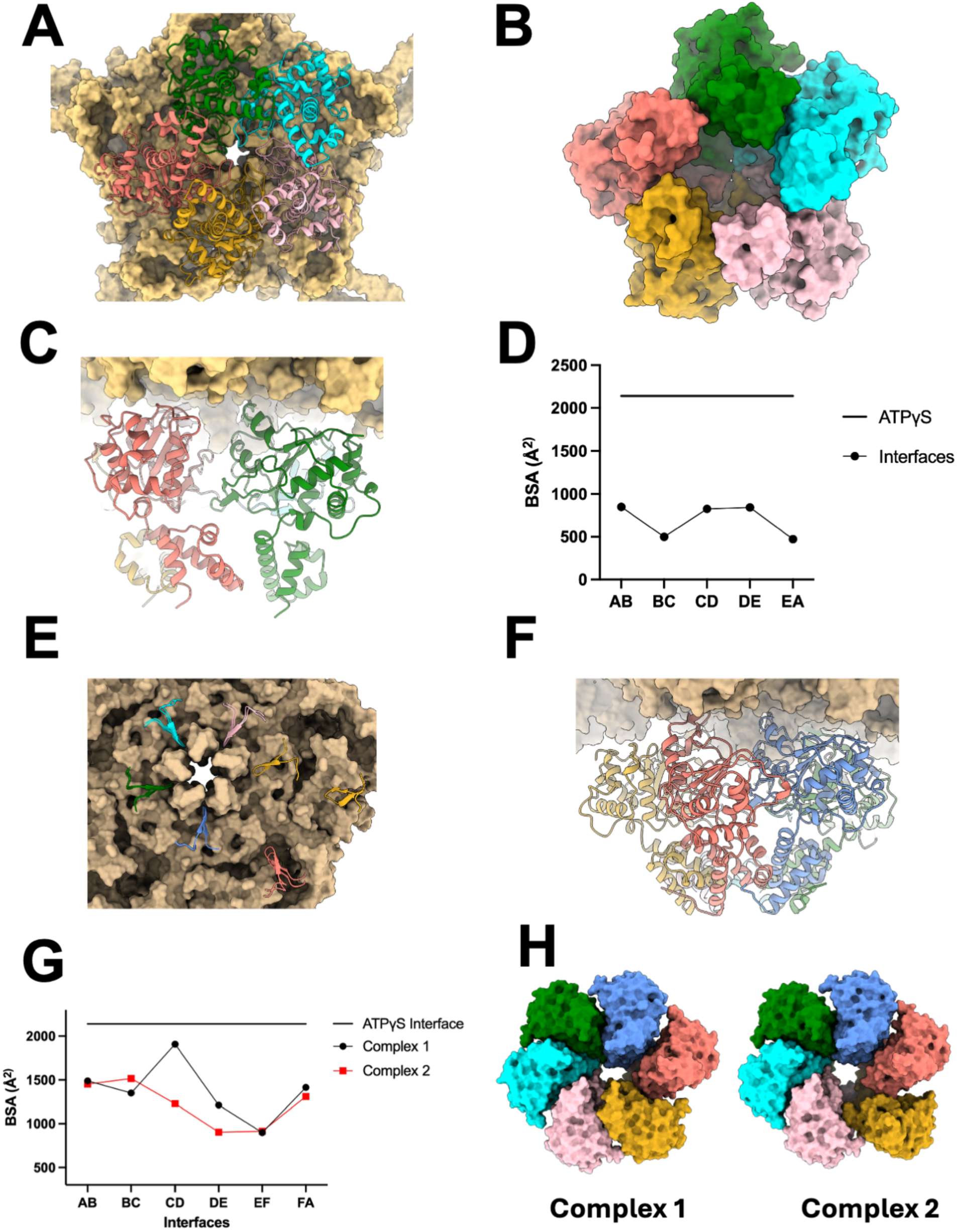
Rep210-Capsid Structures. (A) Rep210 Ribbon representation of the pentameric ring docked into the five-fold capsid pore (Surface representation, tan color). The view is down the fivefold axis. Rep210 chains are colored green, cyan, pink, gold, and salmon for chains A-E, respectively. (B) Surface representation of the docked Rep210 pentameric ring, illustrating that the central channel is closed by the ps1βh loops. (C) Ribbon representation of subunits A (green) and E (salmon) showing the ODs do not interact. (D) Buried Surface Area (BSA) plot of the pentameric ring subunit interfaces, compared to an average ATPγS interface from the Rep68-AAVS1 complex (pdb 9BU7). (E) Ribbon representation of the post-sensor1 β-hairpin2 (pos1βh2) superposition of the two hexameric complexes. The largest differences are in subunit E (gold), where the one in hexameric complex 2, is placed at the edge of the 5-fold VP pentamer. (F) Ribbon representation of hexameric complex 1 subunits, showing the interaction between their ODs, due to the linker latch motif. (G) Buried Surface Area (BSA) plot of the subunit interfaces from the two hexameric ring complexes. The line represents the BSA average from the ATPγS interface of the Rep68-AAVS1 complex (PDB 9BU7). (H) Surface representations of the two Rep210 hexameric ring complexes showing that the inner channel is open.

### Hexameric Complexes

The two hexameric ring complexes dock into the capsid, with only four subunits interacting with the 5-fold pore due to the symmetry breakage between the interacting complexes (Figure 3E). The two undocked subunits remain part of the hexamer primarily due to OD-ring interactions facilitated by the linker-latch residues (Figure 3F). The two complexes superimpose with an RMSD of 0.793 Å, with the most significant difference being the position of one of the undocked Rep subunits, which is farther away from the center of the ring (Figure 3E). We will differentiate the two complexes as tight and loose based on the observed interaction between subunits. Compared to the pentameric complex, the interfaces in the hexamers are significantly stronger, with average BSA values of 1378 Å^2^ and 1221 Å^2^ for the tight and loose complex, respectively. The larger BSA in the tight complex is due to the larger interface between the C and D subunits (Figure 3G). Unlike the pentameric complex, both hexameric complexes feature an open inner channel spacious enough to allow an ssDNA molecule to pass through. The channel dimensions in the tight hexamer at the ps1βh region are approximately 9.6 Å by 22 Å (Figure 3H). At a lower map threshold in the map of the tight complex, we observe density in the inner channel at two locations: at the entrance of the OD ring and in the AAA^+^ ring around the ps1βh loops. This density likely corresponds to ssDNA.

### Post-sensor 1 β-hairpin 2 is the key Rep-capsid interacting motif

Unlike other SF3 helicases such as papilloma virus E1 (PPV-E1) or simian virus T-large antigen (SV40-Tag), AAV Reps are unique because they contain an additional β-hairpin motif (residues 427-434) positioned C-terminal to the ps1βh and aligned parallel to it (Figure 1A). The hairpin contains mostly polar and hydrophobic residues docking into a pocket formed between two pore spikes (DE_1_ and DE_2_) of the narrow 5-fold pore (Figure 3E, 4A). This interaction interface is further stabilized by hydrogen bonds and salt bridges (Figure 4B). Two residues at the tip of pos1βh2, the G430 and N431, make hydrogen bonds with main chain amide Q325 and E322, respectively, both located in spike strand βD_B_ (Figure 4B). In addition, the main-chain carbonyl oxygen of N431 interacts with the main-chain amide NH of T671, located in the HI_B_ Loop. The only direct interactions with the DE_1_ spike are the salt bridge and a hydrogen bond between the Rep residue K385 and D327 (Figure 3B). The main interface interaction is a β-strand expansion, where the second strand of pos1βh2 (βd) induces folding of a region in the VP HI_2_ loop (residues 668-670), forming a three-stranded antiparallel β-sheet between the Rep and VP proteins (Figure 4B-C). The interface is similar in both pentameric and hexameric complexes, with each Rep molecule contacting two VP subunits and covering approximately 800 Å^2^ of surface area; however, more than 2/3 of the interface is between a single Rep and one VP subunit (Figure 4C). The strong VP3-Rep interface prompts us to evaluate whether any AI-based structure-prediction tools (Alphafold3, Boltz2, Chai1) can predict interactions between the Rep helicase domain and VP3. Our results show that only Alphafold3 produced models that closely match the CryoEM Rep-VP3 structure and its interactions (Figure 4D).

**Figure 4.**
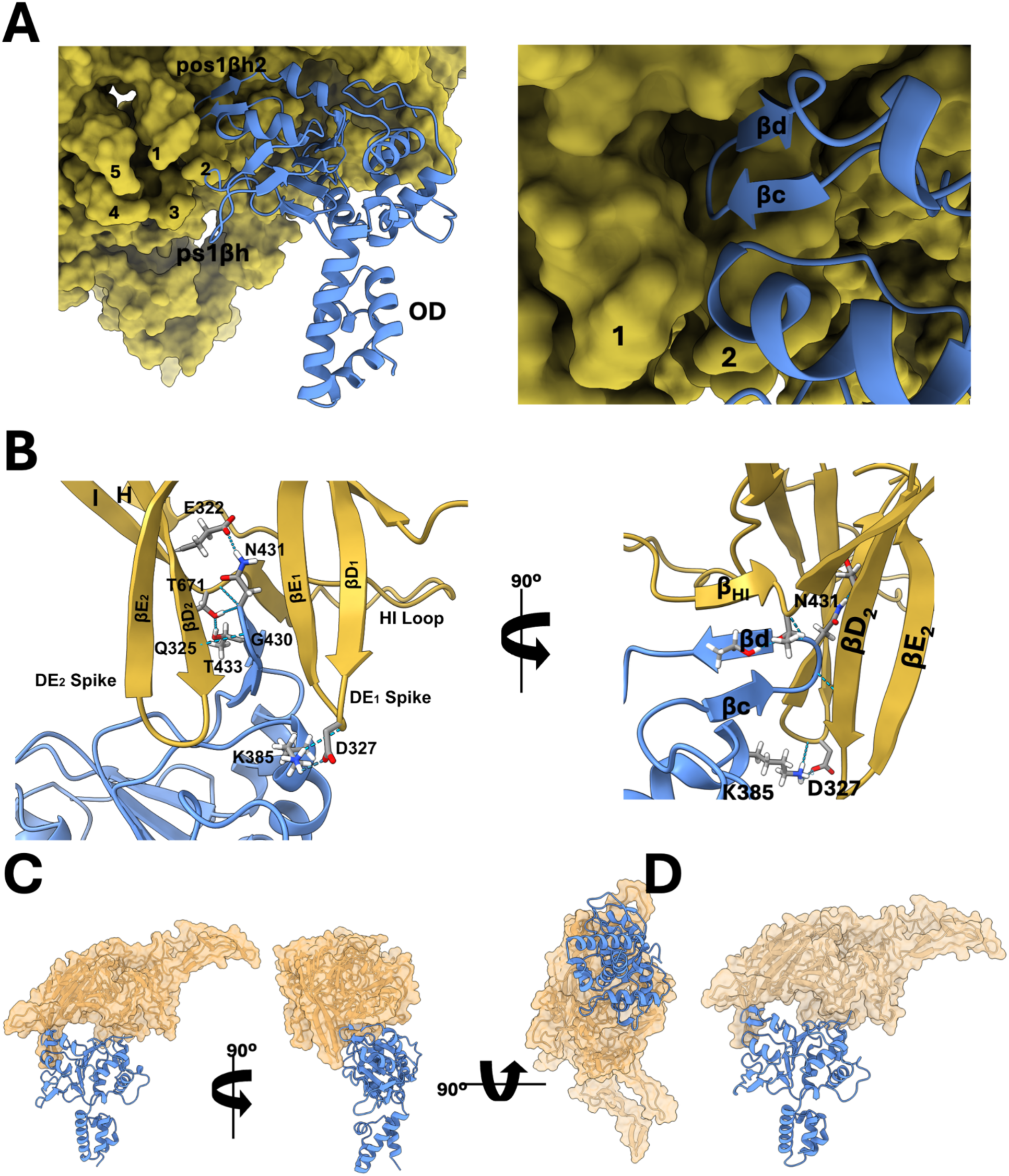
Pos sensor1 β-hairpin2 is the capsid interaction module. (A) Representative interaction between Rep210 (blue ribbon) and the 5-fold pore (tan surface). The numbers represent the five pore spikes. (Right panel) Zoom-in of the Rep-Capsid interaction. (B) Two views of the interactions between Rep210 pos1βh2 residues G430, N431, and pore spike βD residues E322, T671, Q325. There is also interaction between a second spike (D327) and Rep210 K385. Rep pos1βh2 strand β_D_ makes hydrogen bonds with a region in the HI loop inducing the folding of the strand β_HI_. (C) Multiple views of the interaction between a single Rep210 subunit and a single VP3 subunit.

### The poS1βh2 motif allosterically modulates Rep ATPase activity

A key feature of SF3 helicases and Rep proteins is that the interaction of ps1βh with DNA allosterically enhances their ATPase activity^62–64^. Since the poS1βh2 motif is located between strands β4 and β5, we investigated whether poS1βh2 affects Rep ATPase activity (Figure 5A). We created three new Rep68 constructs: Rep68Δβh2, missing the residues from 425 to 438; Rep68VID427-429A, with V427, I428, and D429 replaced by alanine; and Rep68STT432-434A, with S432, T433, and T434 changed to alanine. The mutants were designed using AlphaFold3 (AF3) to confirm that there is no effect on the overall Rep 3D structure. First, all three mutants were tested for their ability to hydrolyze ATP. Figure 5B shows that deleting the pos1βh2 motif greatly reduces its ATP hydrolysis activity to about 8% of WT activity. The ATPase activity of the two alanine mutants is also affected, but to a lesser degree, decreasing to 68% for Rep68STT432-434A and 61% for the Rep68STT432-434A mutant (Figure 5B). In the presence of ssDNA, the activities recovered but never reached WT levels, with mutant Rep68STT432-434A reaching 90% of WT activity. The helicase activity of the three mutants parallels the ATPase results, with the Rep68Δβh2 mutant showing the greatest decrease (40% of WT activity), while the other two mutants show slightly decreased activity (Figure 5C). The results suggest the pos1βh2 may be allosterically connected to the ATPase catalytic center.

**Figure 5.**
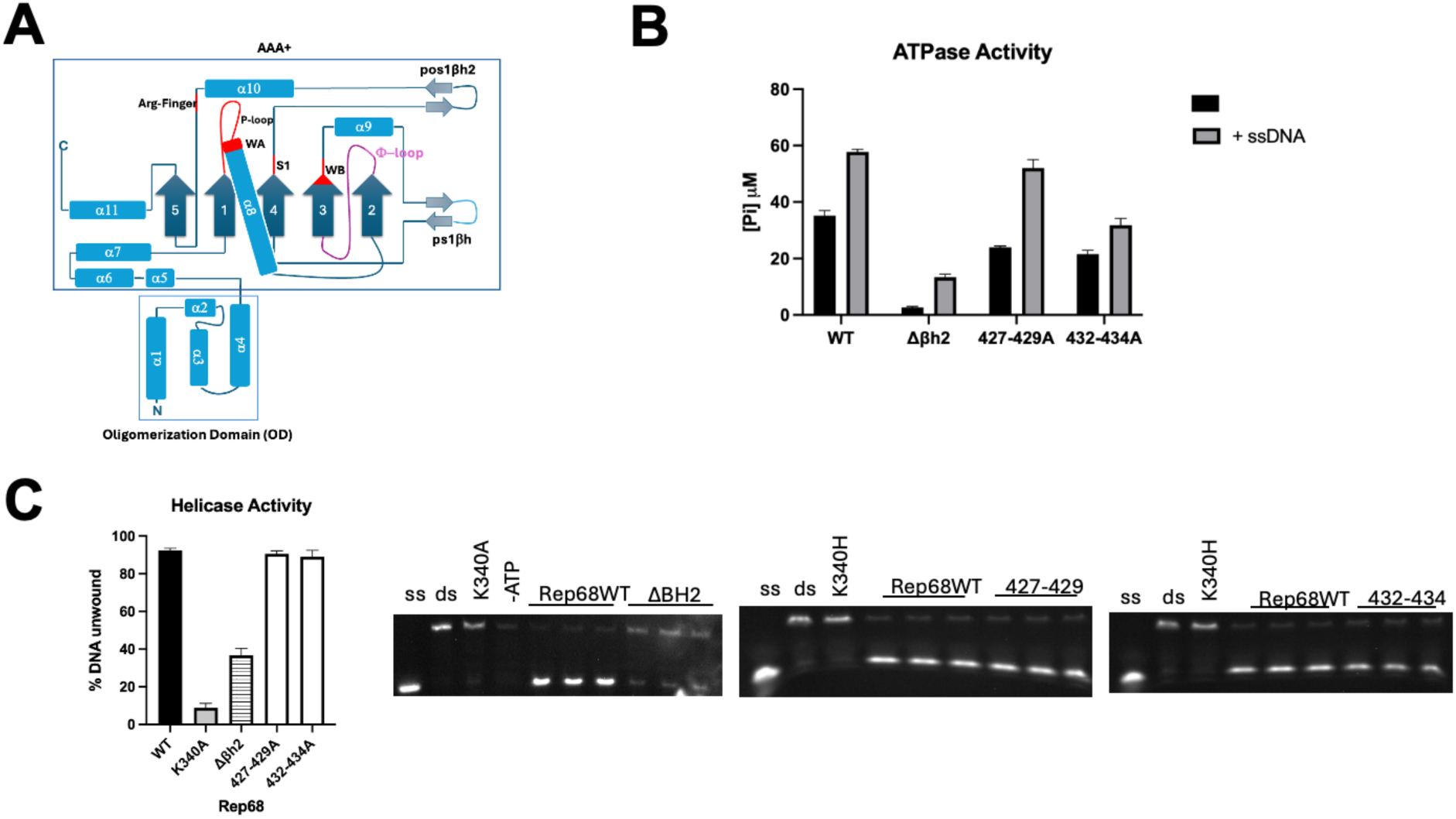
Role of the pos1βh2 motif in Rep catalytic activity. (A) Topology diagram of the Rep helicase domain illustrating that the pos1βh2 is between the sensor 1 and β5, including the arginine finger (R444). (B) ATPase activity of the Rep68 without the pos1βh2 motif (Δβh2), and two mutants with selected residues mutated to alanine (427-429A) and (432-434A). (C) Effect of mutations on the helicase activity. The deletion mutant lost 80% of the wt activity.

### Nucleotide States of the Hexameric complexes

To evaluate the nucleotide states of the hexameric complexes, in addition to analyzing the density in the cryo-EM map, we examined the interface BSA and the conformation of the helicase domains. This was essential when evaluating the presence of ATPγS in cases where the density was ambiguous. In the pentameric complex, all subunits are empty, possibly due to the wide distance between the subunits (BSA_avg_ 849 Å^2^). The two hexameric complexes differ in the nucleotide states of their subunits. In complex 2, the map shows nucleotide density resembling ADP in three subunits (A, B, F), while the other three subunits (C, D, E) are nucleotide-free (Figure 6A). This is represented by the low BSA of the interfaces of the empty subunits, which is the lowest, averaging 1015 Å² (Figure 3G). In addition, the conformations of the helicase domains are ‘open’ compared to the ATPγS-bound conformations seen in previous structures^61,65^ (Figure 6B). The complex 1 map shows density for three ADPs (A, B, F), one empty site in subunit E, and two ATPγS molecules in subunits C and D (Figure 5C). The BSA of the CD interface is the largest, with 1908 Å², comparable to that observed in the Rep ATPγS-bound interfaces (Figure 3G). More importantly, the conformation of subunits C and D is similar to the Rep-ATPγS-bound conformation seen in previous DNA complexes (Figure 6C). Inspection of the subunit C ATPγS molecule shows that important trans residues from subunit D, such as the arginine finger residue R444, K391, and K326, form hydrogen bond contacts, and the interface appears optimal for ATP hydrolysis (Figure 6D). In contrast, the DE interface lacks trans-interactions between subunits, with only R400 interacting with the adenine base and Y354(D) interacting with S407(E) (Figure 6E). This is the result of subunit E without a nucleotide and not having the proper conformation to engage the ATPγS-bound subunit D. All ADP-bound interfaces in the two hexameric complexes (three in complex1 and three in complex2) show no trans-interactions with the ADP molecule.

**Figure 6.**
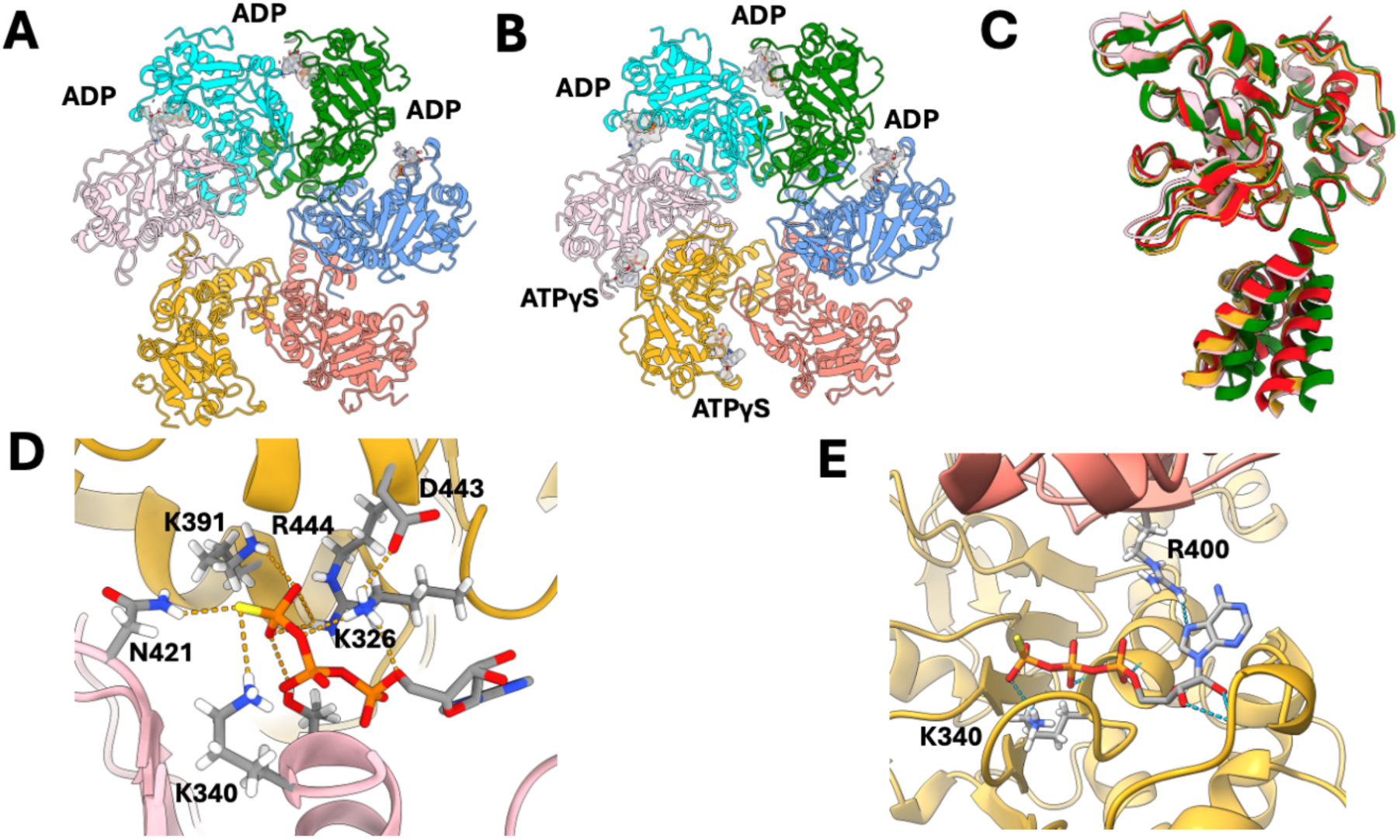
Nucleotide states of the hexameric ring complexes. (A) Ribbon representation of the hexameric complex 1. Re210 subunits A-F (counterclockwise direction). The different bound nucleotides are represented as sticks, and the superimposed Cryo-EM map density for the nucleotide. (B) Ribbon representation of the hexameric complex 2. Re210 subunits A-F (counterclockwise direction). The different bound nucleotides are represented as sticks, and the superimposed Cryo-EM map density for the nucleotide. (C) Superposition of the OD conformational changes elicited by ATPγS binding. Green colored Rep represents a nucleotide-free conformation; Red colored Rep represents the ATPγS conformation; Complex 1 C and D subunits are colored pink and gold, respectively. (D) ATP binding pocket at the complex 1 CD interface. Walker A residue K340 and sensor 1 residue N421 from subunit C, are shown as sticks making hydrogen-bond interactions (orange dashed lines). Representation of subunit D trans-acting residues K391, K326, D443, and arginine finger R444 contacting the ATPγS molecule. (E) ATP binding pocket at the complex 1 DE interface. Walker A residue K340 from subunit D is shown as sticks making hydrogen-bond interactions (orange dashed lines). Representation of subunit E trans-acting residues R400 contacting the ATPγS molecule.

### The structural features of Rep-Capsid interaction are conserved in all AAV serotypes and parvoviruses

The use of AAV cross-packaging systems across different serotypes has been employed over the years as a strategy to enhance packaging efficiency and load capacity, indicating that sequence conservation among serotypes is sufficient for effective cross-Rep-Capsid complex formation and packaging^3,66–70^. Structure and sequence alignment of the AAV1-13 Reps around the pos1βh2 sequence shows that in a stretch of 36 residues, spanning the central β4 to β5 strand regions, there is 100% identity, except for AAV3 and AAV5, where the identity is 97.1% and 94% respectively (Figure 7A). Similarly, the VP sequences in the HI loop segment that interact with Rep, although less conserved than Reps, have an average conserved identity of 84%, with the lowest being AAV5 with 67% (Figure 7B). Significantly, the region within the expanded β-sheet (residues SFI) is fully conserved. Next, using AF3, we asked whether the NS1 helicase domains from other parvoviruses also contain the pos1βh2 motifs. The prediction models indicate that this motif is characteristic of parvoviral helicases, as other SF3 members, such as PPV-E1 and SV40-Tag, lack it (Figure 7B). We then assess whether AF3 can predict similar capsid-NS1 interactions in other parvovirus systems. The models suggest that the interactions and interfaces are likely similar between NS1 and its respective capsids, implying that the packaging complex interaction mechanism could be universal among parvoviruses (Figure 7C).

**Figure 7.**
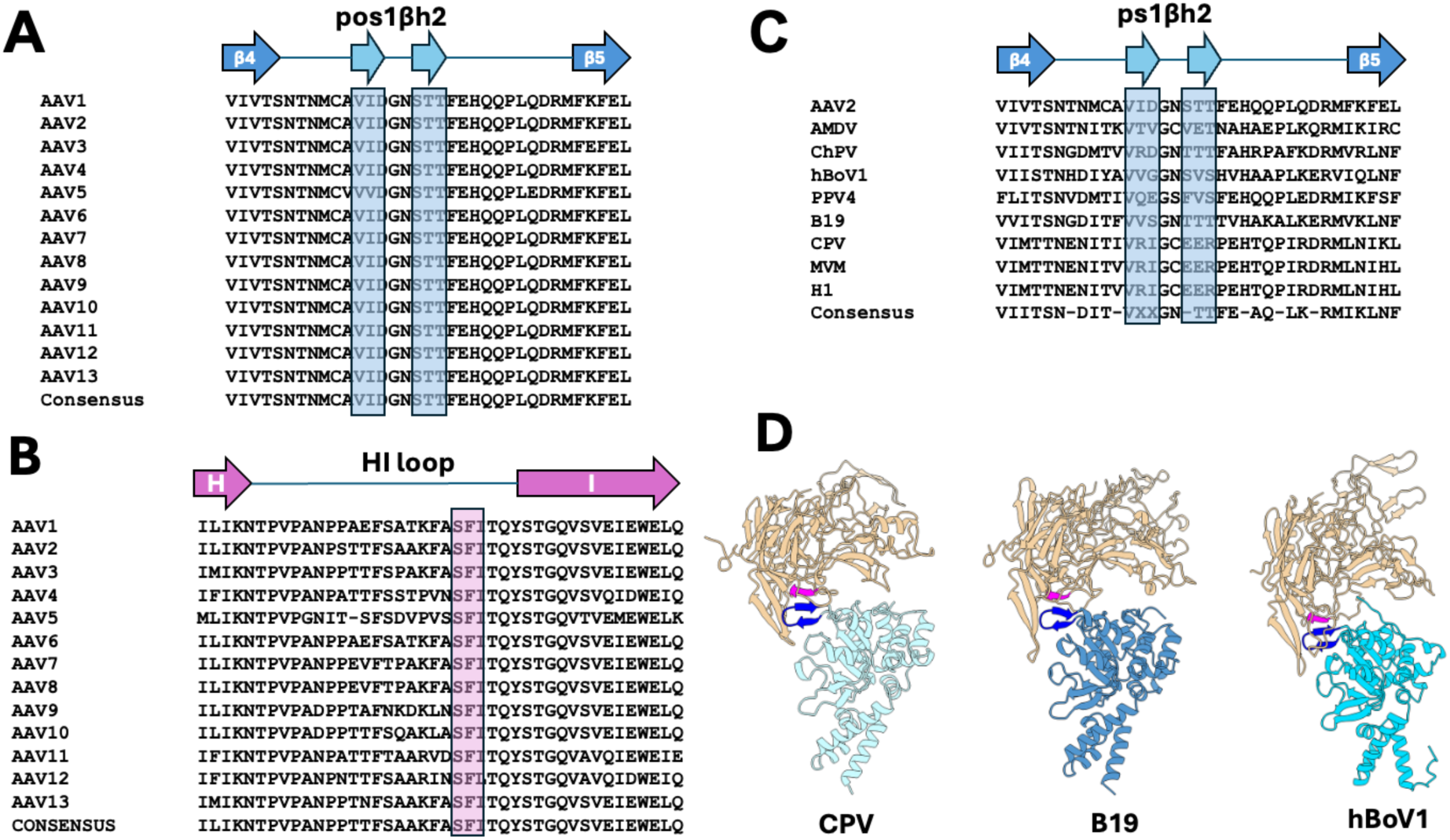
Conservation of capsid interaction motifs among parvovirus Rep and NS1 proteins. (A) Rep sequence and structural alignment between central β-sheets 4 and 5 that include the pos1βh2 between AAV2 and serotypes AAV2-13. Boxed residues are the conserved residues that form the hairpin. (B) VP sequence and structural alignment of the HI loop of AAV2 compared to serotypes AAV2-13 Boxed residues are the conserved residues that interact with the pos1βh2 strand2 and expand the β-sheet. (C) Parvoviral NS1 sequence and structural alignment between central β-sheets 4 and 5 that include the pos1βh2 between AAV2 and Aleutian mink disease parvovirus (AMDV), chipmunk parvovirus (ChPV), human BocaVirus 1 (hBoV1), porcine parvovirus 4 (PPV4), Parvovirus B19 (B19), canine parvovirus (CPV), minute virus of mice (MVM), and H-1 parvovirus (H1). Boxed residues are the residues that form the predicted β-hairpin2. (D) Alphafold3 structure prediction models of parvoviral NS1 helicase domain/VP complex of canine parvovirus (light blue), B19 parvovirus (slate blue) and human bocavirus1 (cyan). Highlighted in dark blue is the pos1βh2, and in purple the equivalent VP HI-loop region that forms the expanded β-sheet.

## Discussion

The results presented here show how Rep proteins recognize and assemble on the AAV capsid and shed light on the packaging mechanism used by ssDNA parvoviruses. The structural features of Rep/Capsid engagement (such as the direct engagement of post-sensor 1 β-hairpin 2 with the capsid’s DE loop) rationalize prior literature, including mutation studies,^71^ and confirm that—as was widely believed already—the fivefold pore is the location of DNA loading. However, since this study does not establish the relevance of individual conformations of Rep oligomers interacting with the capsid, and because capsid-associated Rep210 did not clearly observe the DNA substrate, these structures mark only the beginning of the structural investigation of the DNA translocation process. It is possible that the protein/protein crosslinking used to stabilize the complex trapped one or more “dead-end” conformations, or that protein concentrations and ratios not found in nature led to aberrant structures. In particular, it is difficult to rationalize how the pentameric Rep oligomer would be active in packaging. However, we think the Rep210 complexes shown here represent a Rep68 complex in a ‘helicase mode’. This is backed by earlier research on Rep68-DNA complexes and the Rep68-capsid reconstructions, which are similar to the Rep210-Capsid complexes. (Figure 1F). *In situ* reconstruction of intracellular packaging events will provide one way to establish the relevance of these structures.

Simultaneous engagement of multiple vertices by Rep oligomers suggests there is no induced or pre-existing asymmetry in the capsid itself^47^ that influences the selection of a single packaging vertex. This calls for an alternative explanation for AAV’s strong tendency to incorporate a single, complete genome. One possibility is that the process is under kinetic control, with the time required to insert a unit-length genome into the capsid being short relative to the frequency with which a Rep oligomer finds a capsid in the cytoplasm or the frequency with which a vector genome interacts with a Rep-Cap complex. Another possibility is that when two incompletely packaged genomes collide inside a capsid, one can exit while the other completes translocation. Increasing the ratio of Rep to genome to capsid will improve packaging efficiency in the latter case but may lead to more defective particles in the former case. Future studies to understand how AAV minimizes the formation of mispackaged particles will be important for engineering more efficient in vivo or in vitro packaging.

A common pattern that has emerged in the study of viral packaging systems is the nearly universal need for symmetry mismatch between the portal and the motor complex. In dsDNA bacteriophages such as phi29, the portal is a dodecameric ring, while the motor protein is a pentamer. These features are conserved in other bacteriophages, such as T4, HK97, as well as in herpes simplex virus 1 ^38,42,72^. The 5-fold pore/6-fold motor symmetry mismatch seen in AAV has also been observed in Cystovirus bacteriophage packaging systems, which feature a hexameric motor that attaches to a pentameric pore. This group of enveloped viruses contains a dsRNA genome but packages three (+)-ssRNA segments in sequence^73–76^. Using symmetry mismatch during packaging provides the system with sufficient dynamic flexibility to allow changes in the conformation of the motor subunits, preventing the formation of static structures. This is evident in the case of the Rep pentameric ring docked into the 5-fold pore. The interaction creates a structure with an inner channel that is closed and cannot function as a translocation unit.

Several key aspects of the AAV packaging process remain poorly understood, including the functional oligomeric state of the Rep-ssDNA-Capsid complex and the specific roles of the large and small Reps during packaging. Lessons from dsDNA virus packaging show that their packaging machineries share three key elements: A portal protein that forms a channel for DNA entry at one 5-fold symmetry vertex, a protein that recognizes the viral genome (small terminase), and a motor protein (large terminase) that translocates the DNA. In parvoviruses, there is no dedicated portal protein; instead, a pore formed by five VP proteins at the fivefold vertices is used for ssDNA entry and serves as the platform for Rep recruitment. The small terminase role is played by Rep68/78, which remains covalently attached to the 5’ end of the AAV genome after replication, guiding it to the empty capsids. Furthermore, Rep68/78 also plays the equivalent role of the large terminase since it exhibits motor activity.

In most parvoviruses, a single NS1 protein—equivalent to Rep68/78—is sufficient for genome packaging, and one of the main questions in the AAV packaging mechanism is why small Reps are necessary, even though the large Reps have the same motor domain. King et al. showed that without Rep40/52, there is a significant drop in full-genome packaging and a 20-fold decrease in total genome encapsidation^30^. These findings suggest that Rep68/78 can package DNA, but it is inefficient and requires Rep40/52 to support its motor activity or to complete the packaging process. Since Rep68/78 delivers the genome to the empty capsid, it is reasonable to hypothesize that a Rep68/78 complex at the 3’ ITR initiates genome encapsidation until it reaches a point where the activity of the small Reps is required.

Currently, no kinetic studies exist to inform the mechanistic details of the packaging process, and our Rep-Capsid structures lack the stage at which ssDNA is being packaged. However, together with earlier structures of Rep68-DNA complexes, they provide a basis for proposing a potential mechanism of Rep-mediated DNA packaging. The complex 1 hexamer has two or more of its subunits bound to ATPγS, providing the starting point for the initiation of the ATP hydrolysis and translocation cycle. Subunit D (golden, Figure 8A) provides the arginine finger (R444) that, *in trans*, hydrolyzes ATP on subunit C (pink). During this stage, the ssDNA is bound to the C subunit, but after hydrolysis, it is released and transferred to the D subunit. Next, the empty subunit E (blue) binds an ATP molecule and acts as the new arginine finger donor, hydrolyzing ATP on the adjacent subunit to the left. This cycle repeats around the hexameric ring as ADP molecules are released (Figure 8B). The key step that links ATP binding and hydrolysis to ssDNA translocation is the transition from an empty state to an ATP-bound state. We have previously shown that AAV Rep proteins are distinct from other SF3 helicases in the large conformational change produced upon ATP binding^65,77^. In the Rep-Capsid complex, as the AAA^+^ domain is docked to the capsid, ATP binding results in movement of the OD towards the capsid, pushing the DNA into the 5-fold pore (Figure 6C, 6E). Compared to the nucleotide-free conformation, the OD can undergo a rigid-body rotation as large as 40 degrees. Several residues in the OD, such as R260, K264, and K272, could participate in the DNA translocation mechanism as they line up and point towards the center of the ring (Figure 6D). We and others had previously shown that mutations at R260 and K264 eliminate the production of infectious particles, although the effect may be due to their negative impact on ssDNA replication^65,78^. The proposed model suggests a sequential, coordinated translocation mechanism in which subunits around the ring alternately bind and hydrolyze ATP. This differs from the bacteriophage phi29 dsDNA packaging mechanism, which occurs in two separate phases: a burst in which ATP is hydrolyzed and DNA is translocated, and a dwell phase in which nucleotide exchange occurs^39–41,79–81^. A similar set of nucleotide states was observed in the Rep68-AAVS1 complex. Here, two ATPgS-bound subunits are flanked to the right by a subunit in the process of binding ATPgS, while the subunit to the left was not well defined but appears to be ADP^61^. Single-particle research on the AAV packaging process is essential to fully comprehend this mechanism.

**Figure 8.**
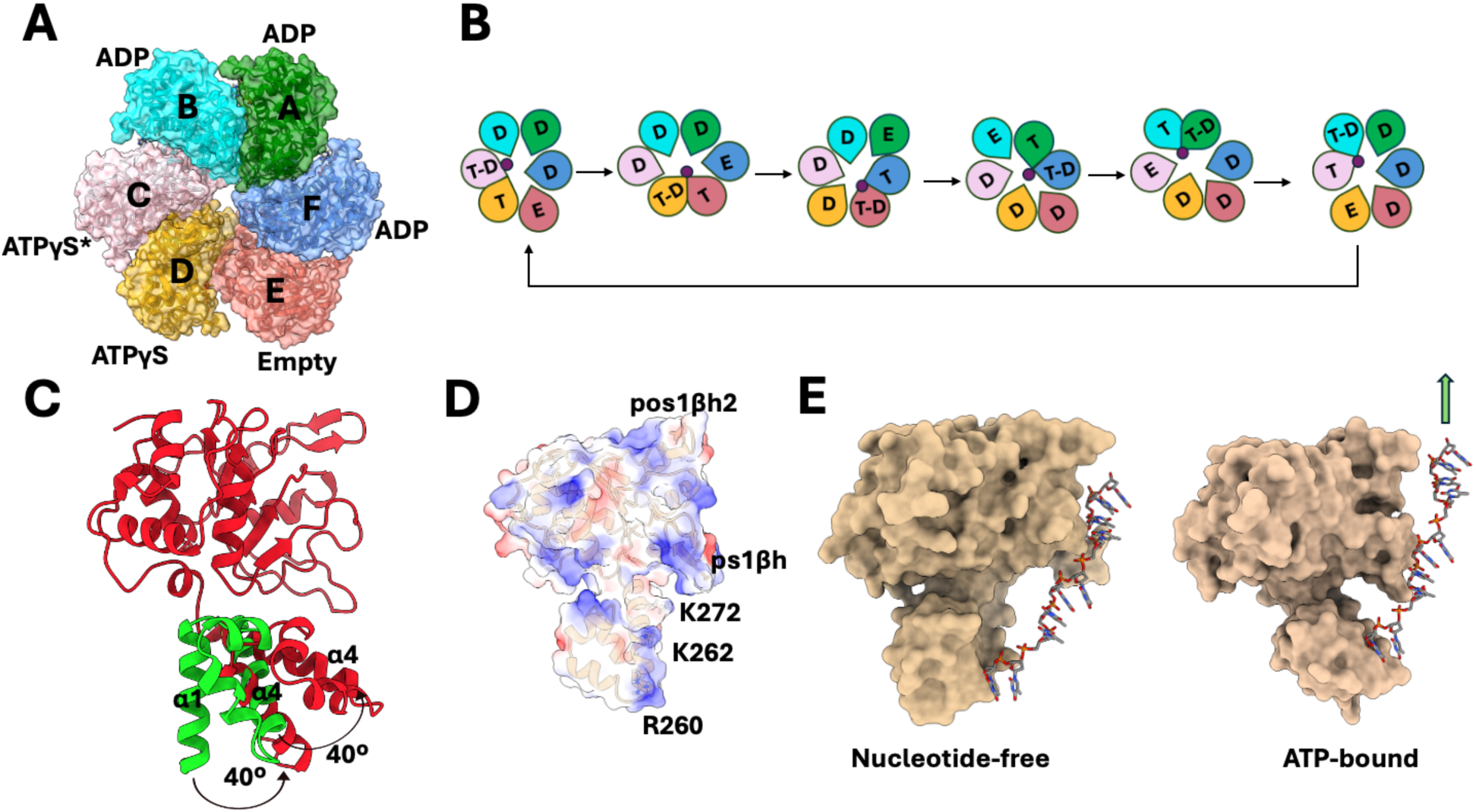
Proposed mechano-chemical cycle coupled to ssDNA translocation. (A) Hexameric ring complex 1 illustrating the different nucleotide states. (B) Proposed mechano-chemical cycle. The different Rep210 subunits are colores as in (A). The dot at the center of the ring represents ssDNA. E = empty state, D= ADP-bound, T-D=ATP-bound, ready for hydrolysis. (C) Superposition of the OD rigid body from nucleotide-free (green) to ATPγS-bound (red). (D) Surface electrostatics of the Rep helicase domain showing the three basic residues that may be involved in ssDNA translocation. (E) Rep surface representation of ssDNA translocation generated by the OD conformational change upon binding ATP.

A question remains regarding the essential function of the small Reps in genome packaging and their role within our proposed model. Since all Rep proteins are likely to interact with the capsid similarly, the key difference between small and large Reps lies in their capacity to form oligomers. While Rep40/52 are monomeric, they may assemble into dynamic hexameric rings with ssDNA and ATP, raising the question of why a Rep40 packaging hexamer might be more efficient than one made by Rep68. One potential explanation is that the Rep40 hexamer can more readily exchange its subunits between ATP-bound and nucleotide-free states. Alternatively, Rep40 may function as a nucleotide-exchange factor aiding in ADP removal. Another possibility is that Rep40/52 could bind to the remaining eleven five-fold pores, as pentamers, thus preventing another Rep68/78 packaging motor from translocating a second genome into the capsid. Further structural and biochemical studies are necessary to resolve these questions. As most prior work dissecting the relative contributions of the large and small Rep proteins relies on deletion or ectopic expression of individual isoforms, it remains unclear whether hetero Rep oligomers play a key role in this process.

A final point from this study is that our results suggest a common packaging mechanism across all parvoviruses, in which the NS1 protein interacts with their capsids via pos1βh2. Given that several studies have explored cross-packaging between AAV and other parvoviruses with mixed results, our results suggest a novel method to create chimeric packaging systems between AAV and other parvoviruses by swapping their pos1βh2 motifs. Ongoing experiments are testing this idea^82–85^.

## Acknowledgements

This study was supported by the National Institute of General Medical Sciences under award number R01GM124204 (to C.R.E.), by a grant from REGENXBIO Inc. (to J.T.K.), and by the National Institute of Allergy and Infectious Diseases under award number R01AI190168 (to J.T.K.). Cryo-EM instrumentation in the Rutgers Cryo-EM & Nanoimaging Facility used in this study was supported by grant S10OD036338, and computing resources were provided by Office of Advanced Research Computing (OARC) at Rutgers, The State University of New Jersey.

## Notes

### Competing Interest Statement

The authors have declared no competing interest.

